# Single-Cell Proteo-Transcriptomic Profiling Uncovers Post-Translational Regulatory Layers in Human Hematopoiesis

**DOI:** 10.64898/2026.10.09.757909

**Authors:** Josephine Gollin, Nieves Garcia-Gisbert, Tomoya Isobe, Elisa Laurenti, Berthold Göttgens, Andrian Yang, Nicola K. Wilson

## Abstract

**Background:** Hematopoietic stem cells (HSCs) have the unique ability to produce all mature cell types in the blood and understanding the mechanisms governing their differentiation and renewal is a driving question in stem cell biology. While single-cell RNA sequencing has transformed our view of human hematopoiesis, mRNA abundance represents an incomplete proxy for functional cell states. Here, we apply single-cell multimodal profiling to simultaneously measure 280 surface and intracellular protein targets alongside whole transcriptomes within individual adult human hematopoietic stem and progenitor cells.

**Results:** By directly coupling modalities within single cells, we show that protein-RNA correlations peak dynamically in transitioning intermediate progenitors across lineages, consistent with pervasive post-transcriptional mechanics underlying lineage choice. Furthermore, key transcription factor proteins maintain multilineage co-expression longer into commitment than their transcripts, demarcating a possible window of fate decision reversibility. Moreover, low and flat *CTNNB1* mRNA levels mask late-erythroid β-catenin protein enrichment, apparently pairing with E-cadherin expression and suggesting post-translational stabilization at the cell membrane.

**Conclusions:** This dataset enables the observation of translational dynamics and RNA-protein relationships throughout the differentiation landscape at an unprecedented level of granularity. Together, our findings demonstrate that transcriptomics alone misses fundamental regulatory programs, establishing single-cell proteo-transcriptomics as a powerful tool for decoding human stem cell fate.

## Introduction

Cell fate decisions made in the bone marrow by hematopoietic stem and progenitor cells (HSPCs) not only maintain steady state blood supply throughout our lifetime, but also dictate our ability to respond to injuries and inflammatory insults throughout life (1,2). Correctly orchestrated, hematopoiesis balances the renewal of hematopoietic stem cells with the generation of committed downstream progenitors and mature cells across hematopoietic lineages (1–3). Understanding the molecular basis of hematopoietic fate decisions has already begun to offer viable pathways towards the modulation of pathogenic hematopoiesis, as exemplified by the success of differentiation therapy in acute promyelocytic leukaemia (4), making it a critical topic of research.

In the era of single-cell transcriptomics, our model of hematopoiesis has shifted from a rigid, tree-like hierarchy toward a dynamic, continuous landscape of fate probabilities (3). Various single-cell studies examined the transcriptome during hematopoiesis to map transcriptional network activity (5,6), building on classical paradigms of transcription factor antagonism—such as the GATA1/PU.1 molecular switch (7). Early models of hematopoiesis established that stem cells express multilineage gene programs prior to commitment (8), and single-cell analysis revealed a spectrum of low-primed hematopoietic stem cell (HSC) states from which more committed progenitors continuously emerge (9). However, surface marker-based sorting and transplantation assays have demonstrated that lineage-biased and functionally distinct progenitors can be isolated at early stages of lineage commitment (10). Moreover, clonally resolved single-cell transcriptomic data have revealed that initial transcriptomic similarity between cells cannot fully explain aspects of cell-intrinsic fate bias (11), suggesting that factors beyond the HSPC transcriptome are critical determinants of downstream fate outcomes during hematopoiesis.

Reconciling these findings requires bridging biological modalities to clarify the role of epigenetic, transcriptomic, and proteomic factors in fate decisions (12,13). Although RNA and corresponding protein expression often positively correlate, correlation can vary widely between individual protein-RNA pairs, demonstrating that mRNA abundance is an imperfect proxy for protein expression (14,15). To precisely link RNA and protein per-cell, <u>c</u>ellular indexing of transcriptomes and <u>e</u>pitopes by <u>seq</u>uencing (CITE-seq) pairs transcriptomic data with surface protein profiles (16) and, applied to hematopoiesis, has allowed for a detailed interrogation of the correspondence between transcriptome and immunophenotype (17,18). However, extracellular markers are often selected for their utility in cell sorting rather than their biological relevance as drivers of differentiation. To gain more insight into the intracellular proteome, where many putative drivers of differentiation reside, CITE-seq has been used to bridge single-cell mass spectrometry of intracellular proteins with the transcriptome (19), a first step towards a more comprehensive description of the intracellular proteome during hematopoiesis. While the detection of low abundance proteins, such as transcription factors, with mass spectrometry remains an active area of development (20,21), the role of such low abundance proteins has typically been imputed from the transcriptome, and the connection between transcription and translation of such proteins remains an open question. To better understand this connection, methods measuring intracellular protein levels and the transcriptome are needed, as evidenced by novel technologies such as SOMA-seq (22).

Here, we apply single-cell multimodal profiling to human hematopoiesis to simultaneously measure intracellular proteins, surface markers, and whole-transcriptome RNA within the same cell. Bypassing the need for inter-modal imputation, this approach provides an direct view into the relationship between the whole transcriptome and the extra- and intracellular proteomes across human HSPCs at an unprecedented scale. Both intracellular and surface proteomic measurements independently recapitulate the hematopoietic differentiation landscape, with protein expression generally mirroring transcript levels while offering a much broader dynamic range. Across the landscape, protein–RNA correlation analysis identified previously unrecognized peaks in transitioning intermediate progenitors and in regions of higher cell turnover. Importantly, we identify notable divergences where transcript levels fail to predict functional protein state, including a novel RNA–protein decoupling predicted to affect human erythropoiesis. Together, these findings establish multimodal proteo-transcriptomics as a powerful tool for decoding human stem cell fate by demonstrating that intracellular protein profiling resolves post-transcriptional regulatory layers invisible to RNA sequencing alone.

## Results

### Combining protein and RNA information yields a rich, multi-omic landscape

We sought to create a comprehensive, single-cell multimodal atlas of hematopoietic stem and progenitor cells (HSPCs) representative of steady state, adult hematopoiesis in order to explore the relationship between the proteome and transcriptome during differentiation. Frozen bone marrow aspirates were obtained from four human donors between the ages of 21 and 33 (Supplementary Table 1). To obtain low-frequency progenitor populations while simultaneously capturing the breadth of cell phenotype necessary to validate antibody staining, we performed magnetic-activated cell sorting (MACS) to enrich CD34^-^ and CD34^-^ bone marrow mononuclear cell fractions, which were subsequently recombined targeting a 60:40 ratio of CD34^+^ and CD34^-^. Cells were sequentially stained with the 75 extracellular and 200 intracellular Multipro antibody-derived tags (ADTs) (Additional Material), as well as with Flex probes against 18,129 genes to generate a multimodal, single-cell landscape (Figure 1A). The final multimodal atlas comprised 137,586 high-quality single-cell profiles. Broad coverage across modalities was confirmed, with a median of 7,755 transcripts and 3,922 unique genes per cell for the RNA modality, and 3,947 counts and 229 unique proteins per cell for the ADT modality.

**Figure 1:**
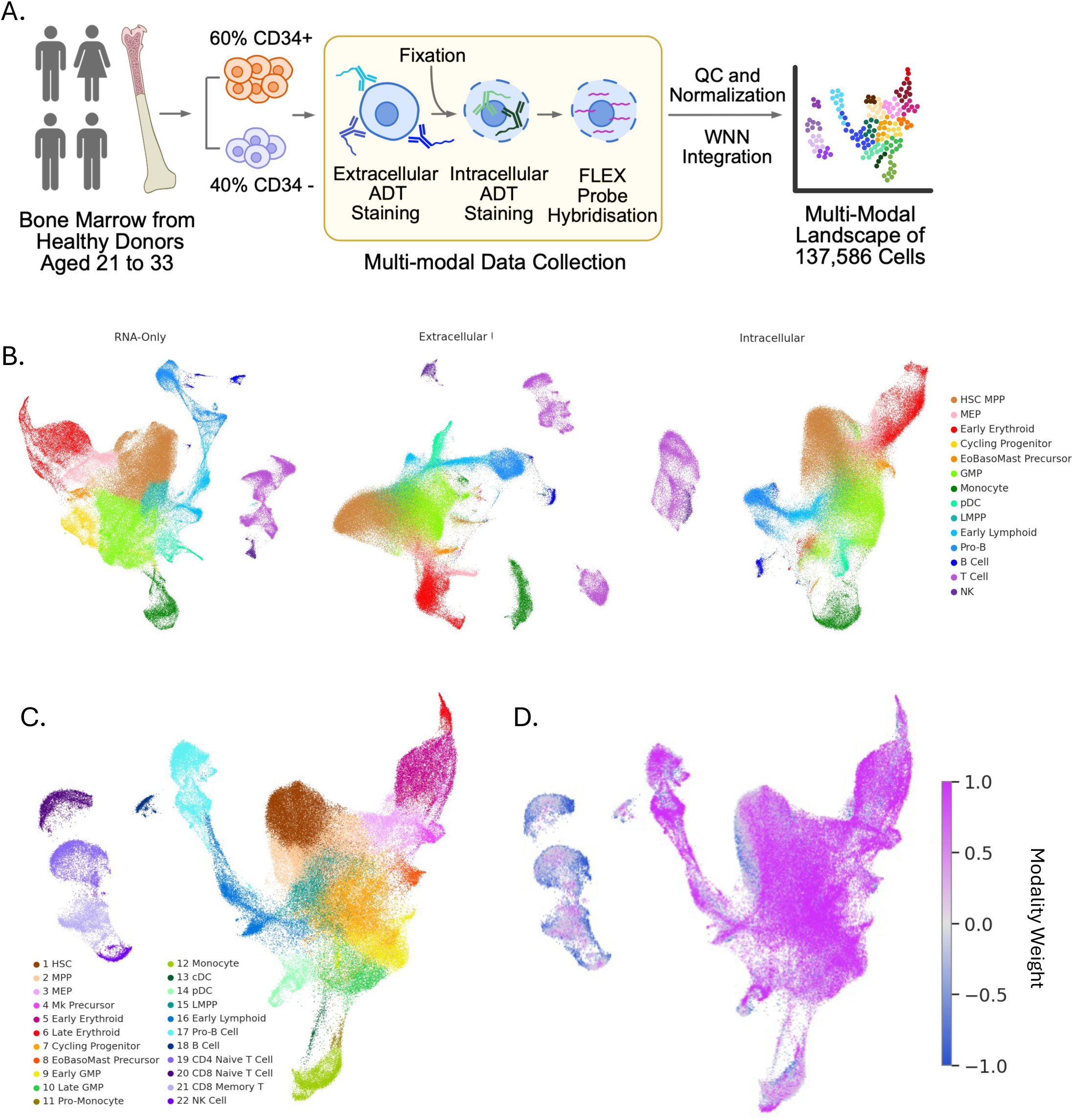
Integration of Protein and RNA information yields an information dense, multi-omic landscape. **A.** Schematic of experimental workflow for Multipro and FLEX data generation and analysis. **B.** UMAP embeddings of the dataset based on, from left to right, only RNA, only extracellular ADT and only intracellular ADT counts, colored based on initial annotations from only the RNA modality. **C.** UMAP embedding based on integration of RNA and ADT modalities via the weighted nearest neighbors algorithm. **D**. Per-cell weights of RNA versus ADT contribution to the integrated graph, with RNA weight as 1 (magenta) and ADT weight as -1 (blue).

Initial annotation of the transcriptome was performed via label transfer analysis using the BoneMarrowMap package (23), and notably, all three modalities were independently able to resolve cell populations defined on the RNA landscape, with similar preservation of the structure of populations and lineages for the RNA, extracellular ADT, and intracellular ADT modalities (Figure 1B). Next, all three modalities were integrated together into a cohesive landscape via the <u>w</u>eighted <u>n</u>earest <u>n</u>eighbors (WNN) algorithm. Final annotation was performed on the joint graph, with label transfer from three RNA-based bone marrow references (18,23,24) and manual refinement using protein information and known marker genes (Figure 1C, Supplemental Figure 1A-C). Cells from each donor were present in all cell type populations (Supplemental Figure 1D). Notably, the ADT modalities contributed more weight to the lymphoid regions of the embedding while the RNA modality contributed more to the early progenitors (Figure 1D, Supplemental Figure 2).

### RNA and ADT modalities mark distinct populations with variable concordance

Across each of the modalities, marker genes and proteins allow the delineation of cell types (Figure 2A-C). However, exclusive marker expression was seen more commonly in the RNA modality, whereas for intracellular and extracellular proteins, populations were distinguished by differences in expression intensity rather than marker exclusivity. RNA and protein expression were moderately positively correlated across all RNA-ADT pairs within the dataset (Figure 2D), with a mean Spearman correlation of 0.18. Extracellular proteins were significantly more positively correlated with their RNAs than intracellular proteins (Figure 2E), which may in part be due to technical differences in the ease of extracellular as opposed to intracellular staining.

**Figure 2:**
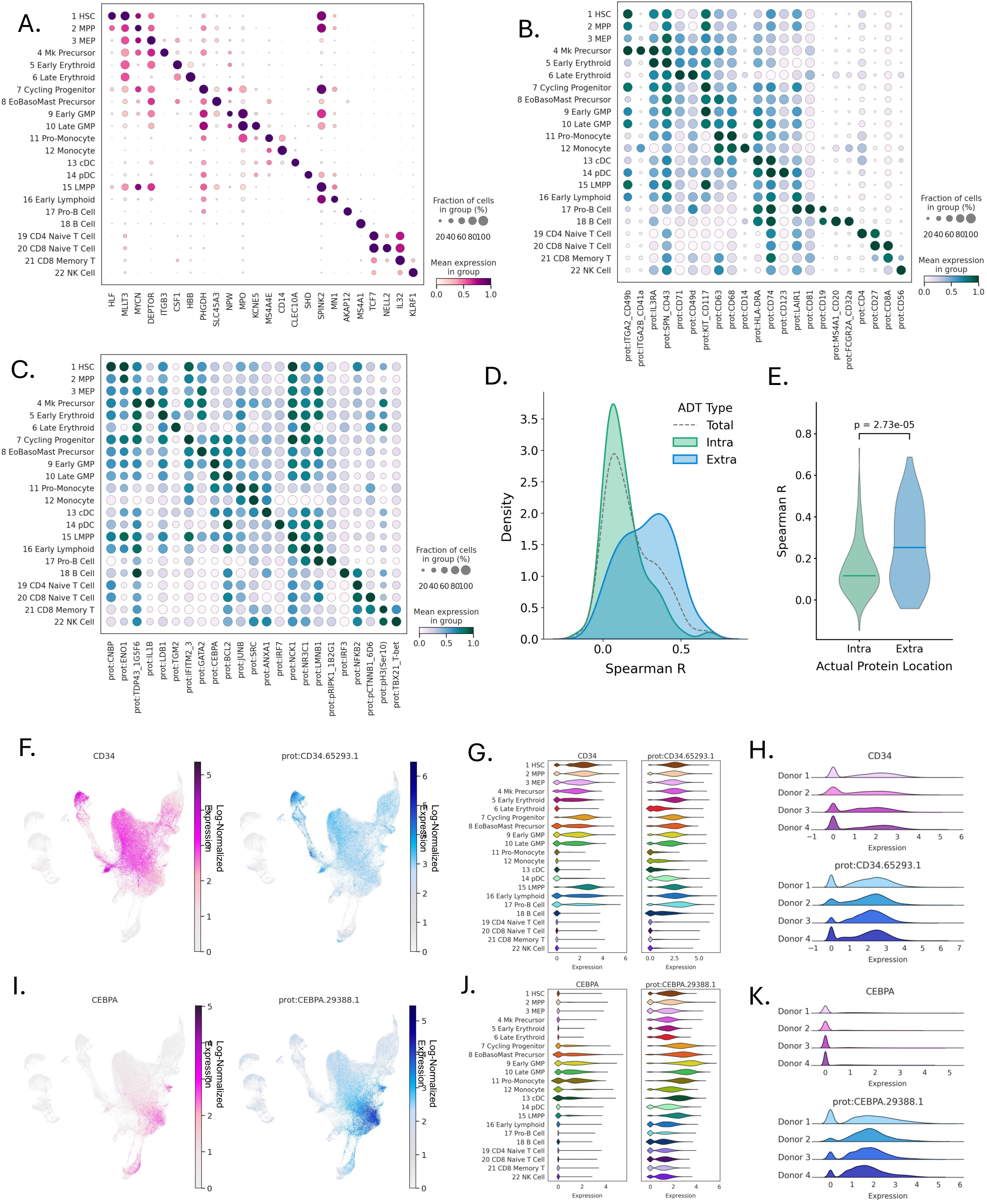
RNA and ADT modalities each mark distinct populations, with positive yet variable correlation between RNA and ADT expression. **A**. Marker genes per cell type B. Extracellular marker proteins per cell type **C.** Intracellular marker proteins per cell type. **D.** Density plot of correlations between intracellular ADTs and RNAs (green), intracellular ADTs and RNAs (blue) and all RNAs and ADTs (grey). **E.** Violin plot of intracellular versus extracellular ADT-RNA correlations. **F.** CD34 RNA (magenta) and ADT (blue) log-normalized expression projected on the UMAP. **G.** CD34 RNA and ADT log-normalized expression per cell type. **H.** CD34 RNA (magenta) and ADT (blue) log-normalized expression per donor. **I.** CEBPA RNA (magenta) and ADT (blue) log-normalized expression projected on the UMAP. **J.** CEBPA RNA and ADT log-normalized expression per cell type. **K.** CEBPA RNA and ADT log-normalized expression per cell type. **H.** CD34 RNA (magenta) and ADT (blue) log-normalized expression per donor.

However, despite the inherent difficulties of intracellular staining, both intracellular and extracellular ADTs captured the same expression gradients as RNA. *CD34* RNA and protein expression occupy the same regions of the landscape (Figure 2F) and are both highest in progenitor cell types (Figure 2G), and *CEBPA* RNA and protein expression both are highest in myeloid progenitor populations (Figure 2I-J). While protein expression generally exhibits a higher baseline expression than RNA, with many cells having non-zero counts of most of proteins assayed, the cell type populations with the highest expression of both were generally concordant (Figure 2G and 2J). Moreover, expression profiles of ADTs were similar between individual donors (Figure 2H and 2K).

### Intermediate progenitor states exhibit elevated RNA–ADT correlation and increased cell to cell variance

Given the power of simultaneous RNA and protein measurements to investigate global protein-RNA relationships during differentiation, we analyzed trends of mean protein-RNA correlation across the landscape. Cells were clustered into neighborhoods using Milo (25) on the joint WNN graph (Figure 3A) in order to facilitate the calculation of correlation metrics in granular and biologically relevant neighborhoods. Diffusion pseudotime and lineage assignment were calculated on the RNA modality alone (Figure 3B, Supplemental Figure 6) with Palantir (26), as lineage inference and pseudotime algorithms based on multimodal graph representations are yet to be developed. Within each Milo neighborhood, we calculated average correlation between all valid RNA-protein pairs, excluding ADTs for phosphorylated epitopes and for multiple protein species. Neighborhoods of intermediate maturity, such as GMPs, appeared to have the highest average protein-RNA concordance (Figure 3C), and along each of the three major lineages inferred from the landscape (erythroid, myeloid, and lymphoid), mean correlation between protein and RNA peaked at intermediate pseudotime (Figure 3D-F). This was observed individually for both the intra and extracellular ADT subsets (Supplemental Figure 7A-H).

**Figure 3:**
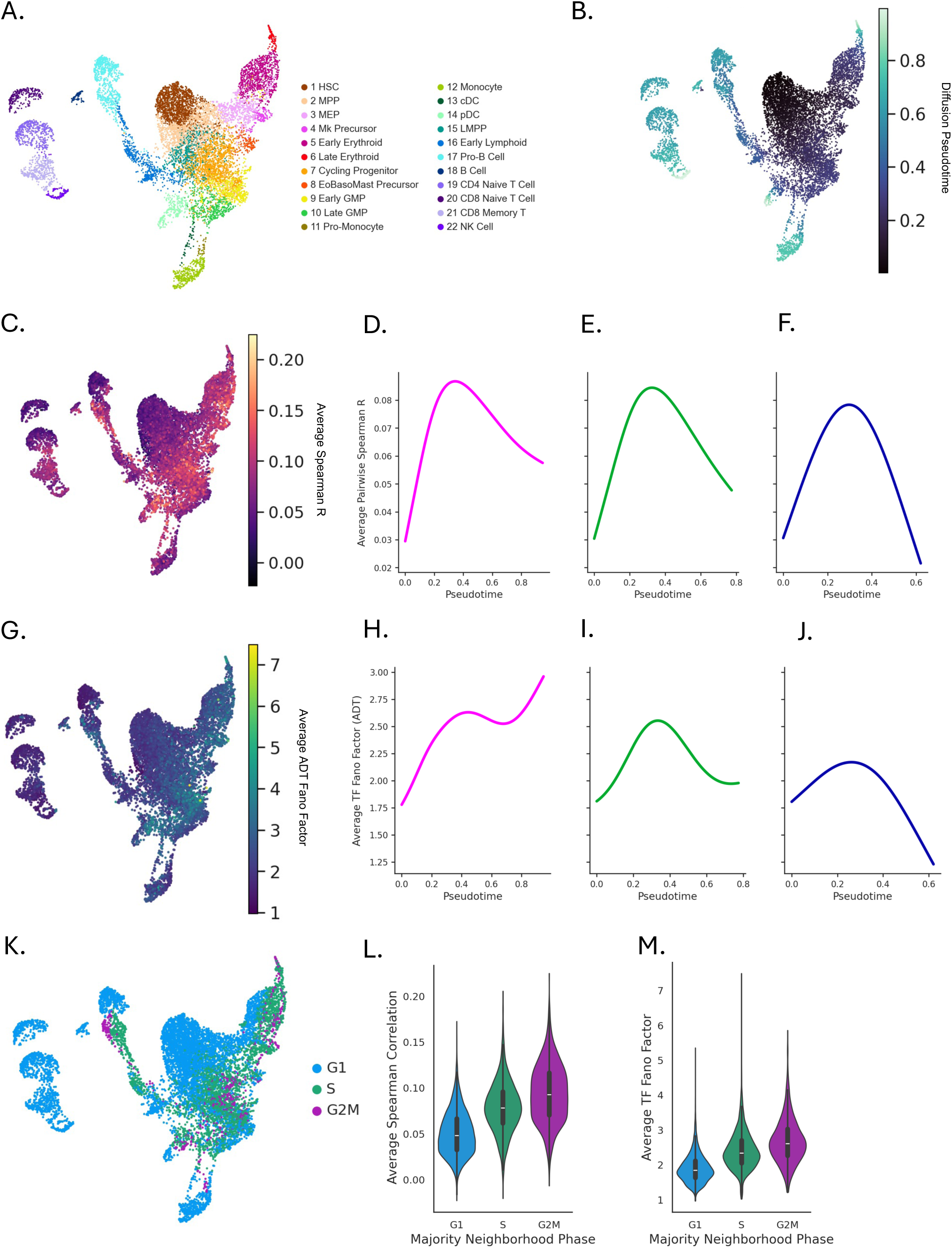
Lineage-independent regions of intermediate maturity have increased intra-cell RNA-ADT correlation and increased inter-cell variability. **A.** Milo-defined neighborhoods projected onto the UMAP, colored by neighborhood majority cell type. **B**. Palantir pseudotime per Milo neighborhood projected onto the UMAP. **C**. Average pairwise RNA-protein Spearman correlation per Milo neighborhood. **D**. Average pairwise RNA-protein Spearman correlation along the erythroid lineage, **E**. myeloid lineage, and **F**. lymphoid lineage. **G**. Average Fano factor of transcription factor ADTs per milo neighborhood. **H**. Average Fano factor of transcription factor ADTs along the erythroid lineage, **I**. myeloid lineage, and **J**. lymphoid lineage. **K**. Majority cell cycle phase per Milo neighborhood **L**. Distributions of mean pairwise Spearman correlation per cell cycle phase. **M**. Distributions of mean Fano factor per cell cycle phase.

Interestingly, interrogating the degree of variability by calculating the Fano factor, a metric used to calculate transcriptional and translational noise (27), across the landscape revealed a similar pattern. The mean Fano factor of the 60 transcription factor ADTs in the Multipro panel (Figure 3G), as well as for their RNA counter parts (Supplemental Figure 7I-L) was highest at intermediate differentiation stages (Figure 3H-J), suggesting that while intracellular correlation between RNA and protein increases, so does cell-to-cell heterogeneity (Supplemental Figure 7M). Finally, assessing the relationship of these trends to cell cycle, we found that in neighborhoods with a high proportion of G2M phase cells (Figure 3K), both mean transcription factor Fano and mean protein-RNA correlation are increased (Figure 3L-M), suggesting a zone of differentiation in which proliferation and molecular turnover increase while the gene regulatory landscape diversifies, likely independent of lineage specification.

### Transcription factor co-expression in progenitors is more apparent in protein than RNA

Given their central role in orchestrating cell fate transitions, we next examined the regulatory dynamics of transcription factors (TFs) across modalities. Both RNA and ADT measurements of transcription factors in the Multipro panel independently recapitulated the global haematopoietic landscape (Figure 4A). Principal component analysis (PCA) of only the transcription factor ADTs was used to rank key ADT TFs driving variation across the dataset (Figure 4B). Despite similar overall explanatory power of protein and RNA expression, a comparison of the top TFs at the RNA and protein level revealed far weaker negative correlations and lineage-specific switches at the protein level than at the RNA level (Figure 4C). A larger dynamic range and lower dropout rate in the ADT modality (compared to the RNA modality), results in smother transition gradients between opposing, lineage defining transcription factors. For example, while at the RNA level, *POU2AF1:GATA2* ratio exhibited a sharp switch, the corresponding ADT protein gradient was much smoother (Figure 4D), with a higher proportion of cells co-expressing low levels of both proteins compared to the RNAs (Figure 4E). This smoothing effect might suggest that transcriptional switches are buffered at the translational level. Moreover, while overall TF protein abundance remained relatively constant across early progenitor transitions, co-expression of combinatorial transcription factor pairs was overall higher at early pseudotime and decreased with maturity (Figure 4F). These findings suggest that lineage-priming programs in early haematopoiesis may be maintained as co-expressed protein networks prior to definitive transcriptional commitment.

**Figure 4:**
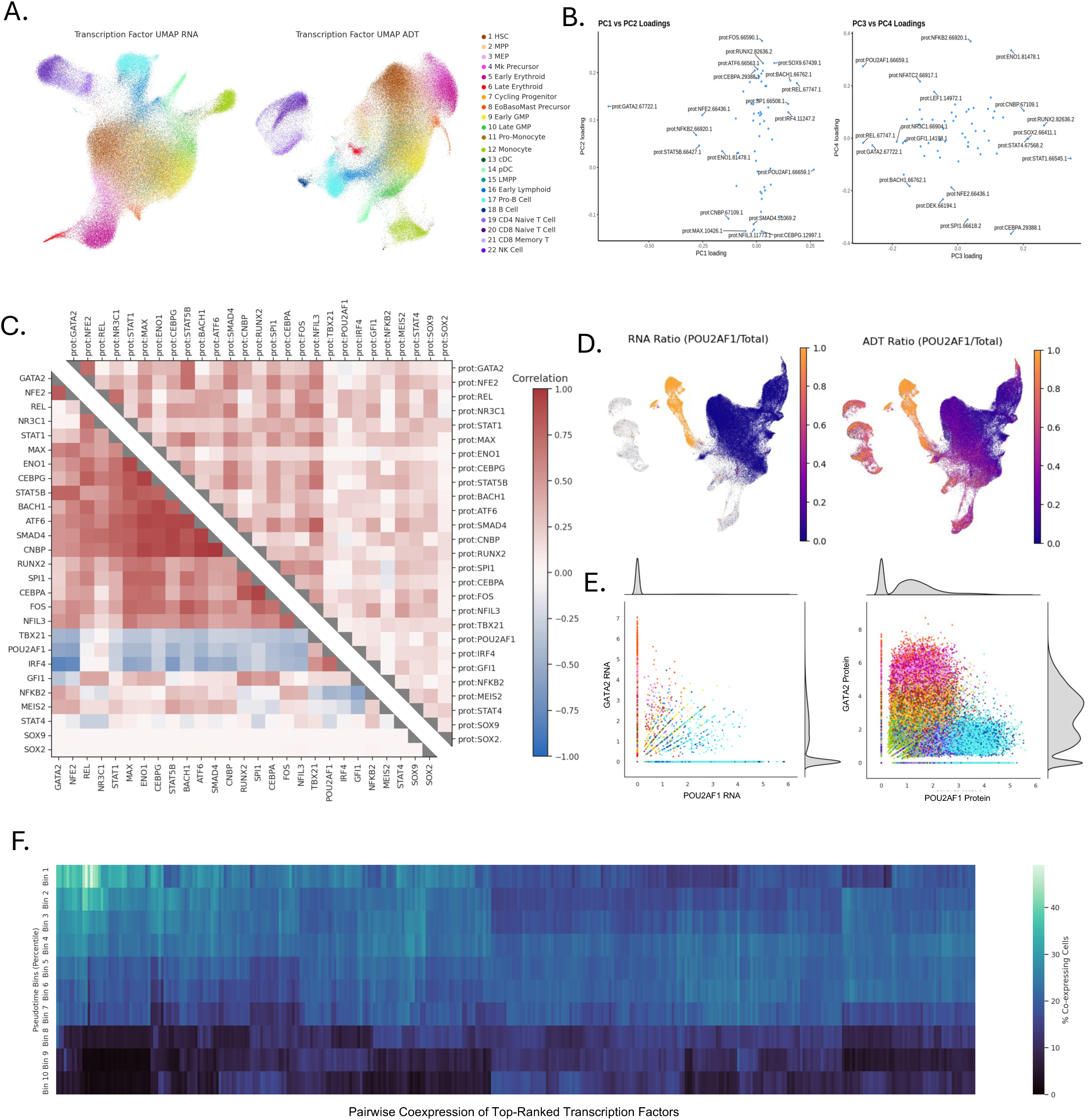
Transcription factor co-expression in progenitors is more apparent in protein than RNA. **A.** UMAP embeddings derived from RNA expression (left) and protein expression (right) of transcription factors in the Multipro panel. **B.** PCA loadings of transcription factor ADTs. **C.** Pairwise correlations and and anticorrelations of RNA (left) and protein (right) transcription factors. **D.** RNA (left) and ADT (right) ratios of POU2AF1 to the total of POU2AF1 and GATA2 expression across the UMAP. **E.** Scatterplots of RNA (left) and ADT (right) expression of POU2AF1 and GATA2. **F.** Heatmap of ADT transcription factor co-expression per pseudotime bin.

### *β*-catenin protein and RNA expression diverge during erythropoiesis

While most proteins exhibited relative concordance with their cognate RNAs, there were notable exceptions. For example, *CTNNB1* (*β*-catenin) mRNA expression was sparse and relatively consistent across the landscape, yet the *β*-catenin protein exhibited marked and dynamic enrichment late in the erythroid lineage (Figure 5A-C). To assess possible reasons for post translational stabilization of CTNNB1, trajectories of highly variable RNAs were matched with that of CTNNB1 along the erythroid lineage (Figure 5D). *WNT* signaling is known to post translationally stabilize CTNNB1 and drive its nuclear localization (28,29); surprisingly, however, canonical components and downstream targets of the *WNT* pathway did not show strong correlation with CTNNB1 protein (Supplemental Figure 8). However, RNA expression of *CDH1* (E-cadherin), an established binding partner of CTNNB1 (30), was tightly correlated with CTNNB1 protein levels along the erythroid trajectory (Figure 5E).

**Figure 5:**
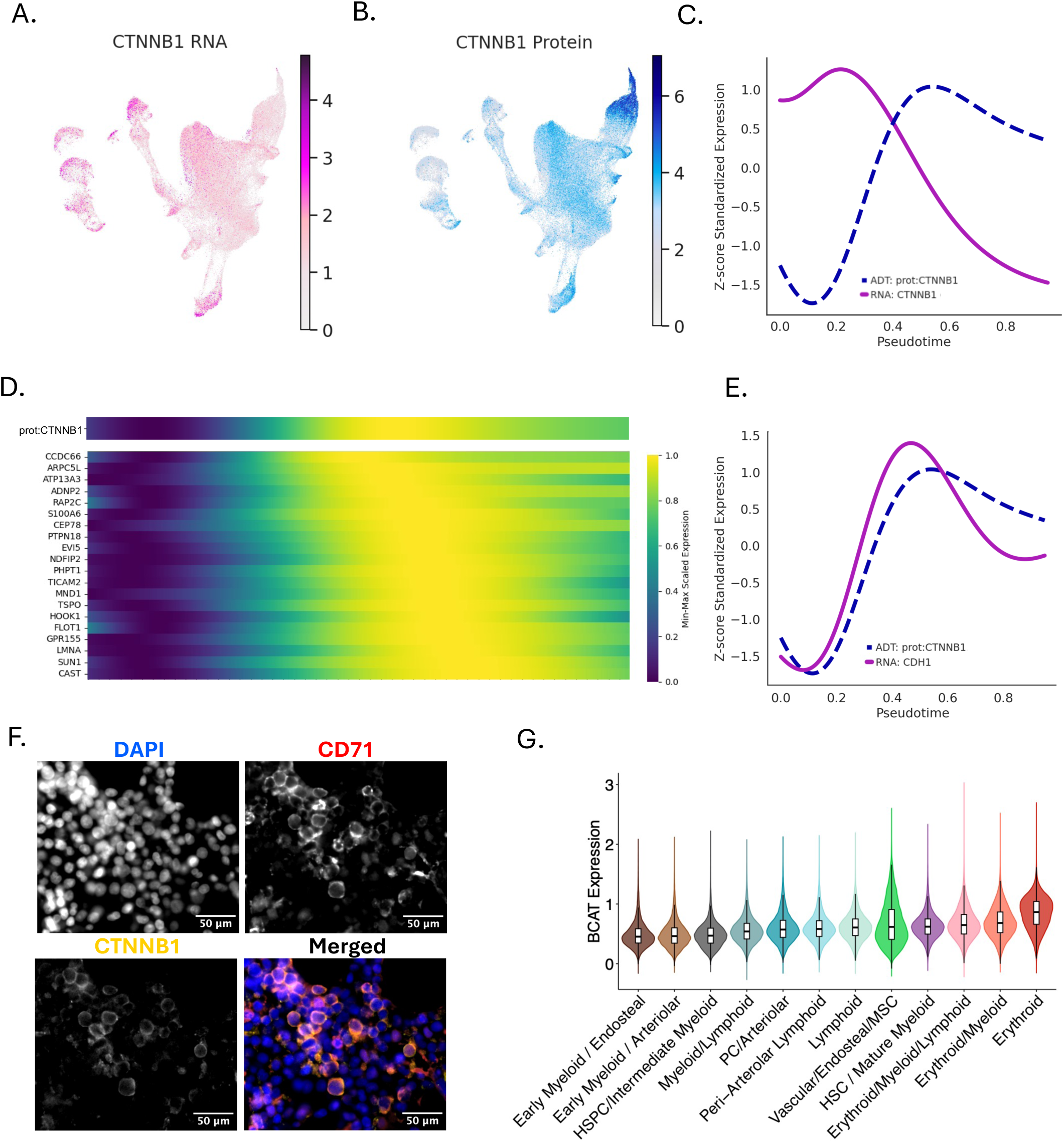
Beta catenin protein expression diverges from RNA in erythropoiesis. **A.** *CTNNB1* RNA expression, and **B.** CTNNB1 protein expression projected onto the UMAP. **C.** Relative *CTNNB1* RNA (magenta) and CTNNB1 protein (blue) expression over the erythroid lineage. **D.** Top correlating RNAs with CTNNB1 protein expression. **E.** Relative *CDH1* RNA (magenta) and CTNNB1 protein (blue) expression over the erythroid lineage. **F.** Representative immunofluorescence images of human bone marrow with nuclear, CD71, and CTNNB1 staining (32). **G.** CTNNB1 protein expression distributions per spatial neighborhood (32).

This tight co-expression matches the established role of E-cadherin in tethering and stabilizing β-catenin at the inner plasma membrane(31), suggesting CTNNB1 may play a structural cell-adhesion role rather than acting as a canonical WNT transcriptional driver during human erythropoiesis. To validate these findings in situ within intact primary tissue, immunofluorescence images of human bone marrow from a published dataset (32) were re-analyzed to confirm robust expression of β-catenin in CD71 positive erythroid regions (Figure 5F), and spatial mapping showed global CTNNB1 enrichment specifically in erythroid neighborhoods (Figure 5G). Crucially, while localized β-catenin protein has been reported (31), all current single-cell transcriptomic atlases have remained completely oblivious to this profound upregulation of β-catenin protein.

## Discussion

In the era of single cell transcriptomics, RNA-based data has been the mainstay of efforts to gain new insights into cell fate decision-making. Given the complexity of post-transcriptional regulatory mechanisms, there are clear limitations associated with reliance on RNA data alone. Investigating the relationship between RNAs and their resulting proteins, particularly for drivers of cell fate like transcription factors, is therefore a worthwhile endeavor. Here, we present a comprehensive, single-cell multimodal landscape of adult human hematopoiesis, evaluating the distinct contributions of the RNA, extracellular, and intracellular protein modalities.

Across cell types, each modality independently resolved major hematopoietic lineages. Integrating them via weighted nearest neighbors (WNN) yields a more cohesive landscape than delivered by any single modality, with surface antibody-derived tags (ADTs) contributing heavily to lymphoid regions, and RNA driving early progenitor mapping. Crucially, because curated antibody panels heavily feature immunophenotypic surface markers, this may reflect composition bias over biology. Extracellular ADTs also displayed tighter concordance with paired transcripts than intracellular ADTs. It remains challenging to decouple whether this disparity stems from biological differences in the flow of the central dogma between secretory and cytosolic proteins, or from technical artifacts introduced during cell fixation and permeabilization. Nevertheless, distinct modalities reveal complementary biological features. While single-cell RNA detection exhibits classic binary dropout, protein quantification distinguishes populations through continuous signal intensity. This may reflect both technical factors in antibody staining but also fundamental biology, including longer protein half-lives relative to mRNA turnover (33,34).

Despite these technical distinctions, multimodal profiling uncovers previously unrecognized dynamics within lineage-uncommitted progenitors. For example, cells at intermediate stages of differentiation exhibit both elevated intra-cellular RNA–ADT correlation and increased cell-to-cell variability (Fano factor). This suggests a dynamic state where high transcriptional and translational turnover coincides with multilineage diversification. While our results do not explain whether and how these molecular features may control cell fate decision processes, it is intriguing that this observation is not limited to a single lineage but instead represents a dynamic process that is observed across lineage trajectories, consistent with a role in broader mechanistic aspects of cell fate decision making. Furthermore, transcription factors demonstrate low-level co-expression at the protein level until later fate commitment. This stands in sharp contrast to the much earlier divergence of RNA expression, primarily used for previous inference of mechanisms (8,35,36). One attractive speculation is that this divergence demarcates a critical window of fate choice initiation versus commitment, wherein prolonged protein persistence maintains a permissive regulatory state in which cell fate decisions remain plastic.

Importantly, our multimodal approach uncovers profound post-translational regulation that remains completely invisible to single-cell RNA sequencing, exemplified by the divergence of β-catenin protein and transcript dynamics. In classical WNT signaling, β-catenin activity is governed not by transcriptional induction of CTNNB1, but by post-translational evasion of the GSK3β destruction complex (29,37). Moreover, β-catenin executes dual cellular functions: acting either as a nuclear transcription factor (canonical WNT) or as a structural adapter tethered to E-cadherin (CDH1) at adherens junctions (non-canonical structural role) (28,31,38). This dual functionality and post-translational control are well-documented in other biological systems, including epithelial-to-mesenchymal transition (EMT) (39), embryonic stem cell pluripotency niches (40–42), and colorectal tumorigenesis (43,44), where membrane E-cadherin acts as a stoichiometric “sink” that sequesters β-catenin to regulate cell adhesion independently of nuclear WNT target activation. In human erythropoiesis, we similarly observe that CTNNB1 mRNA levels remain flat and uninformative, while β-catenin protein accumulates and tightly correlates with CDH1 expression, consistent with a structural, membrane-tethered role rather than a canonical WNT transcriptional program. Because single-cell RNA atlases rely strictly on CTNNB1 transcript abundance, prior studies have consistently overlooked post-translational dynamics in hematopoiesis, underscoring the necessity of co-measuring intracellular proteins.

A key caveat of antibody-based assays is their semi-quantitative nature. Unlike mass spectrometry, ADTs rely on binding affinities that can vary widely between antibodies, making absolute quantification and comparative abundance difficult. Given the role of protein concentration in differentiation (45,46), quantitative proteomics are necessary to gain a full understanding of the role of the proteome in hematopoiesis, but high-throughput antibody profiling serves as a vital bridge, establishing a clear and high resolution mapping from transcription to translation, deepening our understanding of cell-state transitions.

## Conclusions

Here, we showcase a novel methodology to measure intra- and extracellular proteins along with transcripts within single cells in the context of steady-state adult hematopoiesis. We expose previously unrecognized post-transcriptional regulatory layers, including dynamic changes in RNA-to-protein correlation along differentiation trajectories as well as pinpointing where non-canonical WNT signaling operates during human erythropoiesis. Taken together, our findings demonstrate how mRNA abundance is an incomplete measure of stem cell states and differentiation processes, thus establishing single-cell proteo-transcriptomics as a powerful approach for decoding the molecular drivers of human cell fate.

## Materials and Methods

### Sample Acquisition

Bone marrow aspirates from deceased organ donors were harvested at the Cambridge University Hospitals NHS Trust Addenbrooke’s Hospital by the in accordance with regulated procedures approved by the relevant Research and Ethics Committees and with appropriate consent (REC 15/EE/1052 and REC 07/MRE05/44 respectively) by the Cambridge Biorepository for Translational Medicine (CBTM). Bone marrow aspirates from living donors were obtained from Stem Cell Technologies. All processing of human samples was performed in accordance to REC ethical approval (REC IDs: 15/EE/0152 and 24/EE/0116, Cambridge Blood and Stem Cell Biobank study).

### Generation of MultiPro Data

Cells were thawed in a 37°C water bath and mixed with 10 mL Rich Thawing Media (RTM) dropwise, then spun for 5 min at 500 x g, and resuspended in 1 mL PBS/FCS (3% FCS by volume, PBS from Gibco and FCS from PAN-Biotech). The cells were then counted and spun down for 5 min at 500 x g, then resuspended with 90 µL PBS/FCS, 15 µL FcR (Miltenyi) and 15 µL CD34-conjugated magnetic beads (Miltenyi) per 1x108 cells. The cells were then left to incubate for 30 minutes at room temperature, then washed with 10 mL PBS/FCS. The cells were then spun down and resuspended in 2 mL MACS buffer (Miltenyi) and passed through a cell strainer (Greiner), then run through a primed LS column (Miltenyi) according to manufacturer’s instructions. The cells were then stained according to *Proteintech MultiPro Individual Antibody and Cocktail Staining* Protocol (C10011), with the following modification, fluorescently labelled antibodies (CD34-APCCy7 (Clone 581, Biolegend), CD19-FITC (Clone HIB19, Biolegend) and Zombie Aqua (Biolegend)) were added simultaneously with Human TruStain^TM^ FcX buffer and before the addition of extracellular Multipro cocktail. The cells were incubated on ice for 10 mins. The cell suspension was then prepared for FACS, where all live cells were sorted into 0.04% BSA solution (Ambion). Samples were then stained with lyophilized pre-fixation antibody cocktail, and washed once. The samples were then fixed for 18 hours according to the 10x *Genomics Fixation of Cells & Nuclei for Chromium Fixed RNA Profiling* protocol, then frozen and stored at -80°C. The samples were then thawed, Flex probes were hybridized along with antibody barcodes for 24 hours at 42°C, cells were counted and library prep was performed according to the *Chromium Fixed RNA Profiling Reagent Kits for Multiplexed Samples with Feature Barcode technology for Protein using Barcode Oligo Capture* protocol from 10x Genomics. Samples were sequenced on an Illumina Novaseq X.

### Processing of CITE-seq Data

After sequencing, reads were aligned to the human genome (GRCh38, 2020A) and count matrices were generated using Cell Ranger (Version 8.0.1). Quality control on called cells was performed by discarding cells with high levels of ADT dropouts as well as cells with unusually high numbers of isotype controls with the *DropletUtils* (47) package. Quality of cells was also assessed via the percentage of mitochondrial RNA reads, the total RNA and ADT reads, and the diversity of RNA and ADT reads detected. RNA reads were normalized for cell size using a cluster-based approach to estimate cell size with the *calculateSumFactors* function from the *scran* (48) package in R and normalized reads were log transformed. ADT reads were normalized by the geometric mean then log transformed.

### Data Integration and Annotation

Batch correction and integration was performed on both modalities separately using the *FastMNN* function in the *batchelor* (48) package in R. The RNA and ADT modalities were integrated using weighted nearest neighbor analysis (WNN) from the *Seurat* package in R(49). Cells were then annotated based on label transfer from a human bone marrow scRNAseq atlas (23) using the *BoneMarrowMap* package in R (23) , as well as label transfer from two additional atlases (18,24) using *Azimuth* (49). Initial cell types were assigned as a consensus of majority cell types per Leiden cluster, and initial clusters were refined based on marker genes and proteins. Marker genes were curated through both manual selection and through the *FindMarkers* function in *Seurat* (49).

### Metacell Construction and Pseudotime Calculation

Metacells were constructed based on the joint protein-RNA wknn graph generated via WNN analysis using the *makeNhoods* function from the *MiloR* package(25). This resulted in 11,154 neighborhoods. Diffusion pseudotime was calculated solely on the PCA space of the RNA modality using the results of fastMNN integration with the *Palantir* package(26), as *Palantir* was not developed for use with multimodal graph embeddings. Trajectory inference was also performed with *Palantir* based on the RNA modality, selecting terminal states in three major lineages.

### Correlation and Fano Factor Calculations

To calculate correlations between protein-RNA pairs across cells, the Spearman correlation of each RNA and respective protein, excluding phosphorylated ADTs and ADTs against multiple proteins within a family, was calculated per donor and averaged to produce a mean Spearman correlation per pair across all cells. In order to situate trends on the hematopoietic landscape, protein-RNA correlation was also calculated per Milo-defined neighborhood, and the Fano Factor for the 60 transcription factors included in the panel was calculated per neighborhood as the variance to mean ratio of raw counts for the RNA and ADT modalities independently.

### Transcription Factor Selection

In order to select transcription factors at the ADT level that explained a large proportion of the variance, fastMNN was run on ADT counts of only transcription factors. PCA loadings were calculated from the corrected PCA space and 27 transcriptions factors contributing highly to variance in the first 5 PCs were chosen as a subset of highly explanatory transcription factors. To calculate ADT-ADT correlations and corresponding RNA-RNA correlations for these transcription factors across cells, pairwise Spearman correlations were calculated between RNAs and ADTs of interest using vectorized rank transformation.

## Declarations

### Ethics approval and consent to participate

Human samples were harvested in accordance with regulated procedures approved by the relevant Research and Ethics Committees (REC 15/EE/1052 and REC 07/MRE05/44 respectively). All processing of human samples was performed in accordance with REC ethical approval (REC IDs: 15/EE/0152 and 24/EE/0116, Cambridge Blood and Stem Cell Biobank study).

## Consent for publication

Not applicable.

## Availability of data and materials

All data generated in this study will be available via GEO shortly. Code used to process data, create figures, and analyze ADT-RNA correlation will be available closer to publication.

Proteomic imaging data was accessed here: https://plus.figshare.com/collections/Mapping_the_Cellular_Biogeography_of_Human_Bone_M arrow_Niches_Using_Proteomic_Imaging/7174914.

## Authors’ contributions

Conceptualization, N.K.W, B.G. E.L., methodology, N.G.G. and N.W., formal analysis, J.G, A.Y. T.I., investigation, J.G., N.G.G., N.K.W., software, J.G., A.Y., T.I., visualization, J.G., supervision, N.K.W, A.Y., B.G., E.L., writing—original draft, J.G., writing—review and editing, J.G, N.G.G., T.I., E.L., B.G., A.Y., N.K.W., project administration, N.G.G., N.K.W., E.L., B.G., funding acquisition, B.G, N.K.W., E.L.

## Acknowledgments

The authors would like to thank Jason Huang and Kit Nazor for their assistance in developing and implementing the Multipro panel. They also thank Krishnaa Mahbubani and the staff from the Cambridge Biorepository for Translational Medicine for assistance in sample acquisition and Reiner Schulte and the Cambridge Institute for Medical Research Flow Cytometry Core Facility for assistance in flow cytometry. The authors gratefully acknowledge the Cambridge Stem Cell Institute (CSCI) Genomics Facility for their support and assistance in this work.

---

**Supplemental Figure 1:**
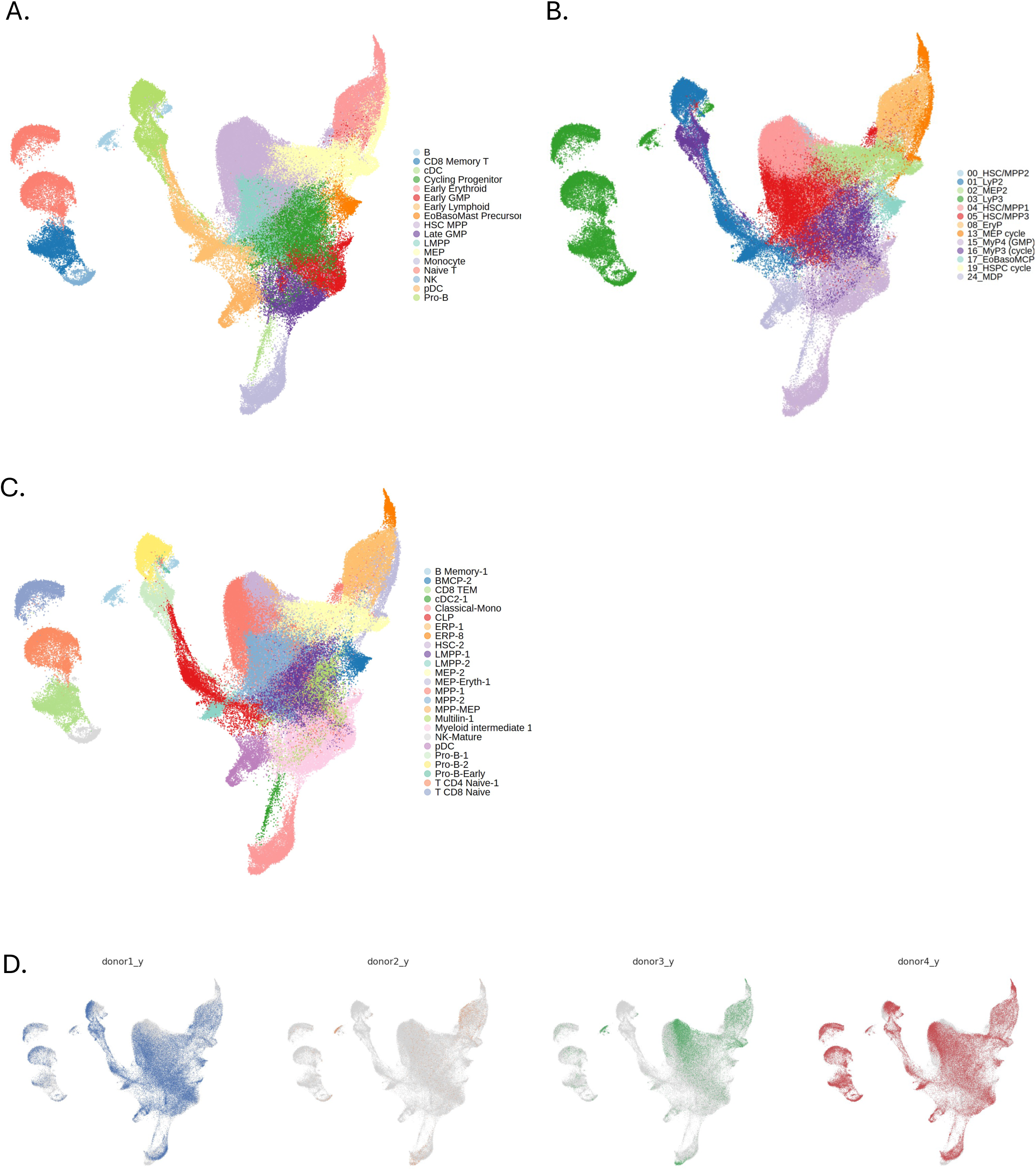
RNA-based label transfer from transcriptomic atlases onto multimodal landscape. **A**. Majority cell type per Leiden cluster based on label transfer from Zeng *et al. .* **B.** Majority cell type per Leiden cluster based on label transfer from Mende *et al. .* **C.** . Majority cell type per Leiden cluster based on label transfer from Zhang *et al.* **D.** Landscape with cells per donor highlighted.

**Supplemental Figure 2:**
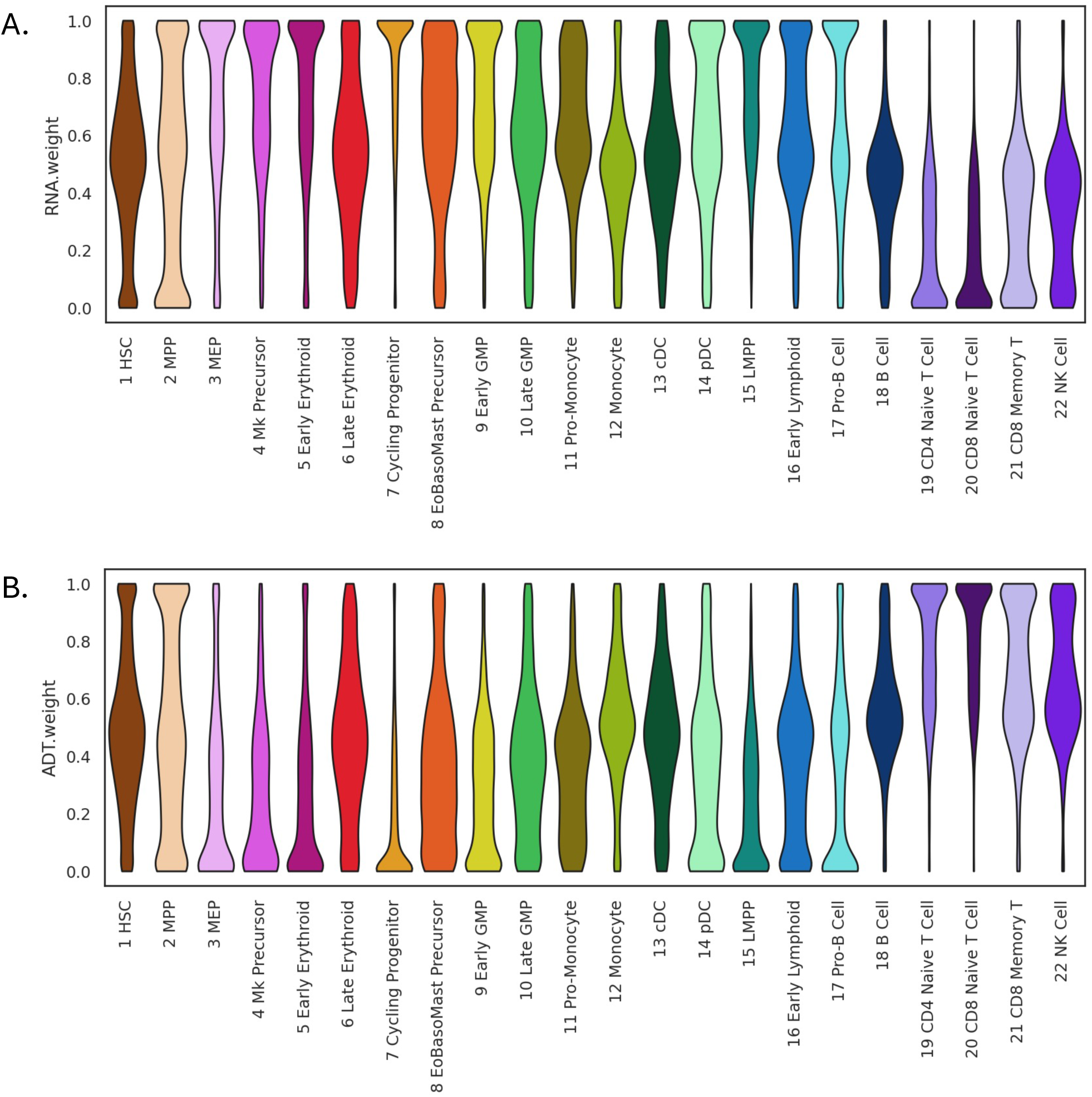
RNA and ADT expression profiles have differing contributions to cell populations. **A**. Violin plots per cell type of RNA weight per cell in weighted nearest neighbors graph construction. **B.** Violin plots per cell type of ADT weight per cell in weighted nearest neighbors graph construction.

**Supplemental Figure 3:**
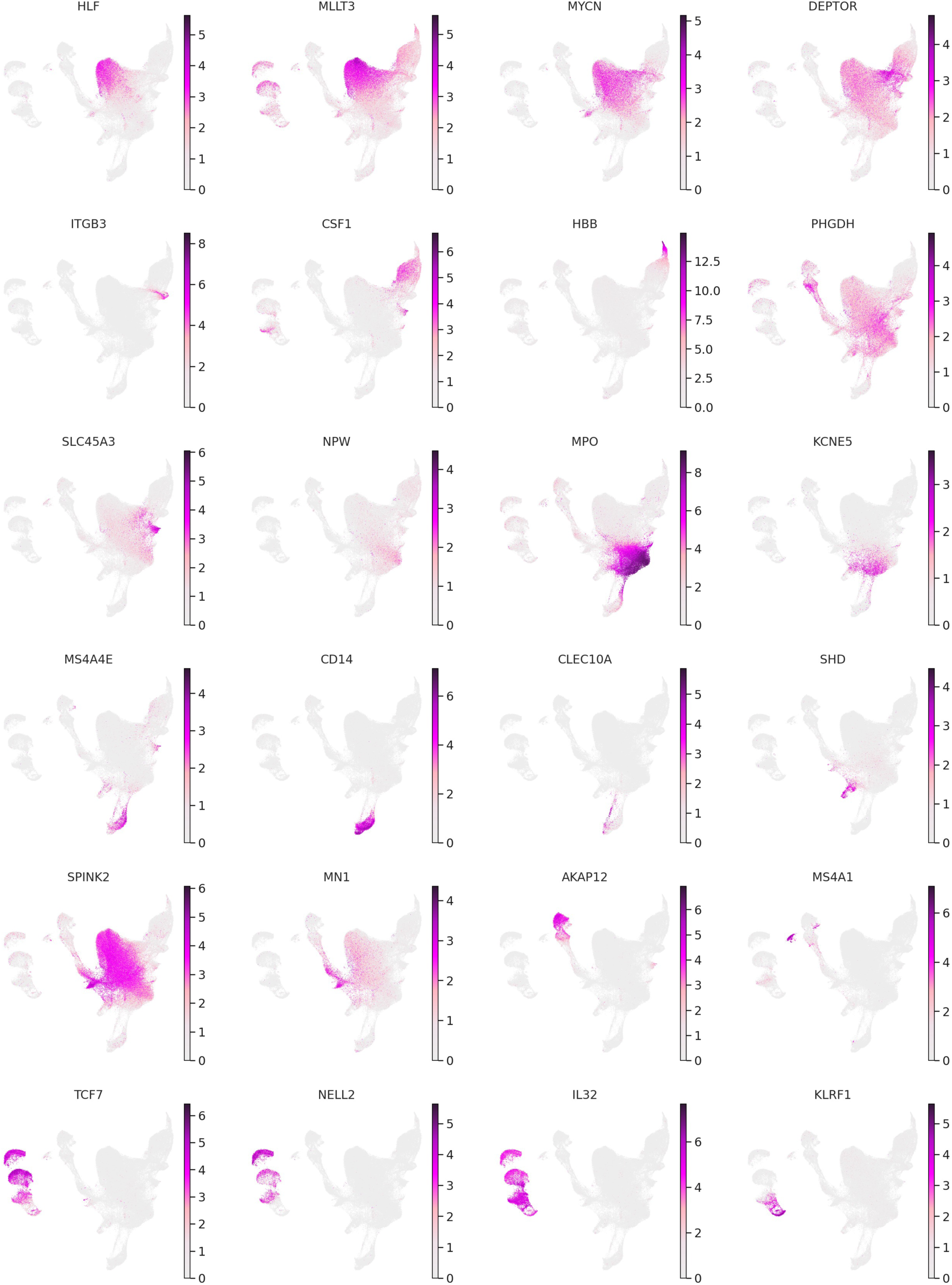
Marker genes projected onto the UMAP. Color represents log-normalized RNA expression.

**Supplemental Figure 4:**
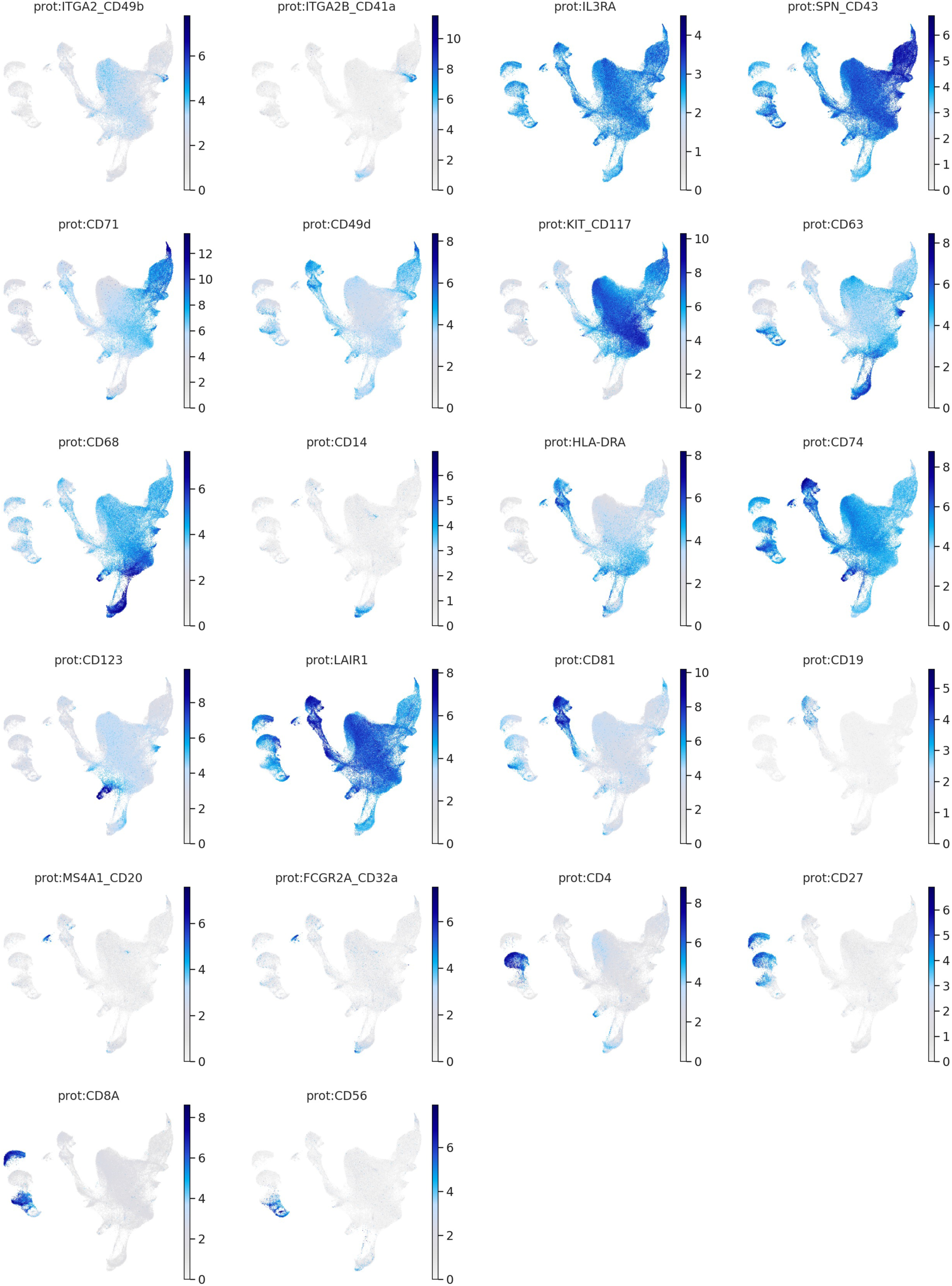
Marker extracellular proteins projected on the UMAP. Color represents log-normalized ADT expression.

**Supplemental Figure 5:**
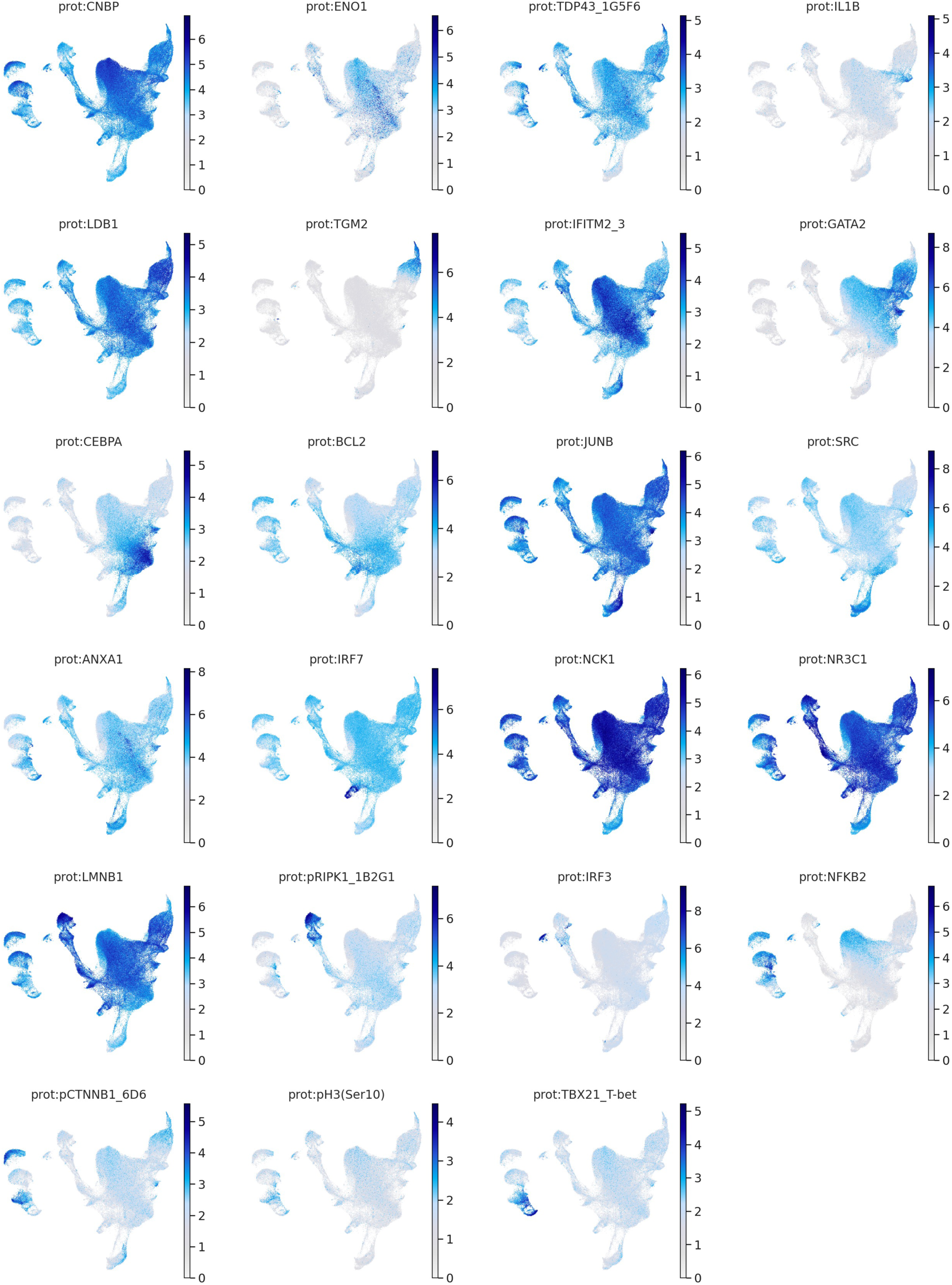
Marker intracellular proteins projected on the UMAP. Color represents log-normalized ADT expression.

**Supplemental Figure 6:**
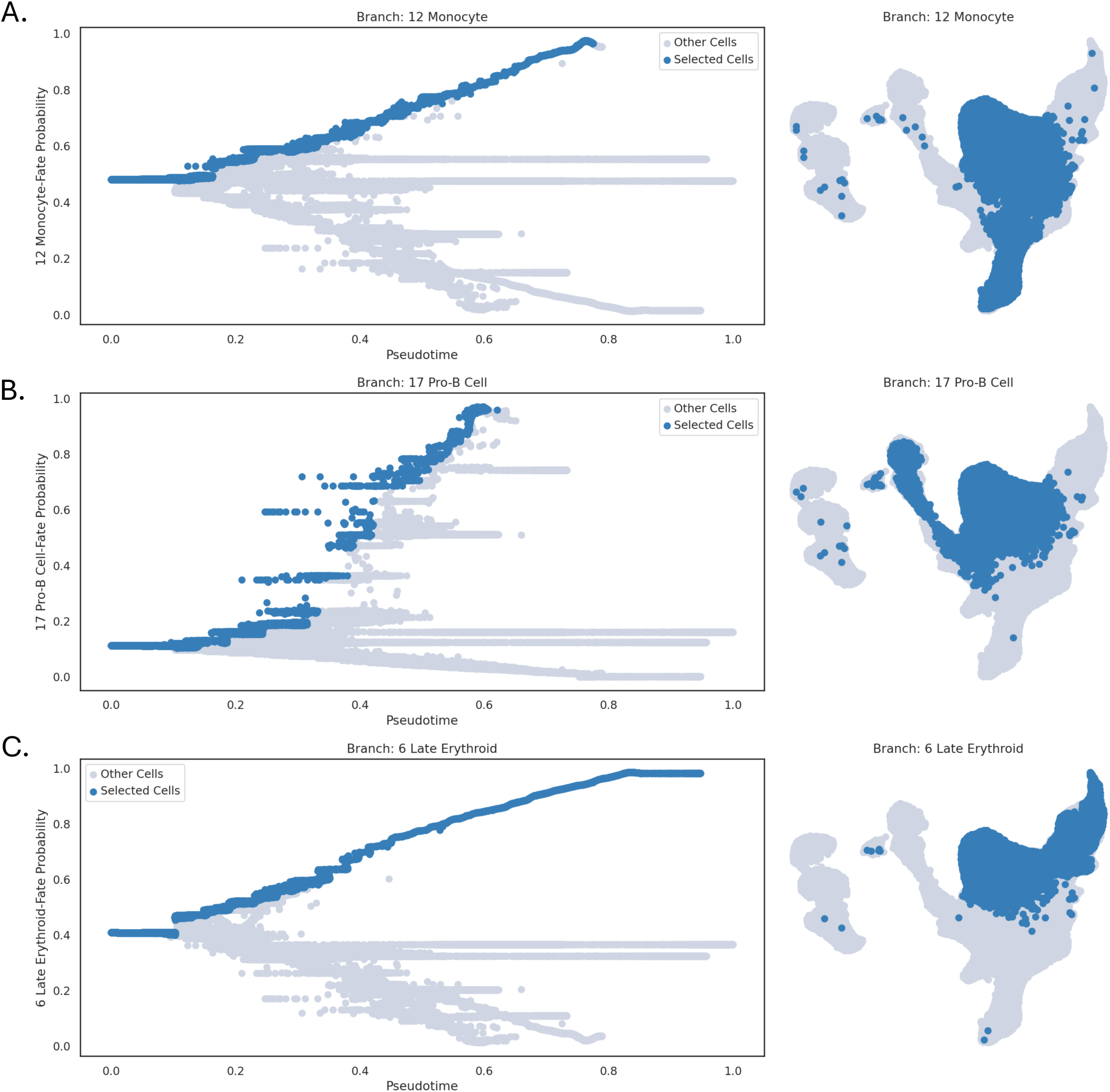
Major haematopoietic lineages defined via Palantir trajectory inference. Fate probability per cell (left) and thresholded cells per lineage (right) for the **A.** myeloid (Monocyte), **B.** lymphoid (Pro-B Cell), and **C.** erythroid (Late Erythroid) lineages.

**Supplemental Figure 7:**
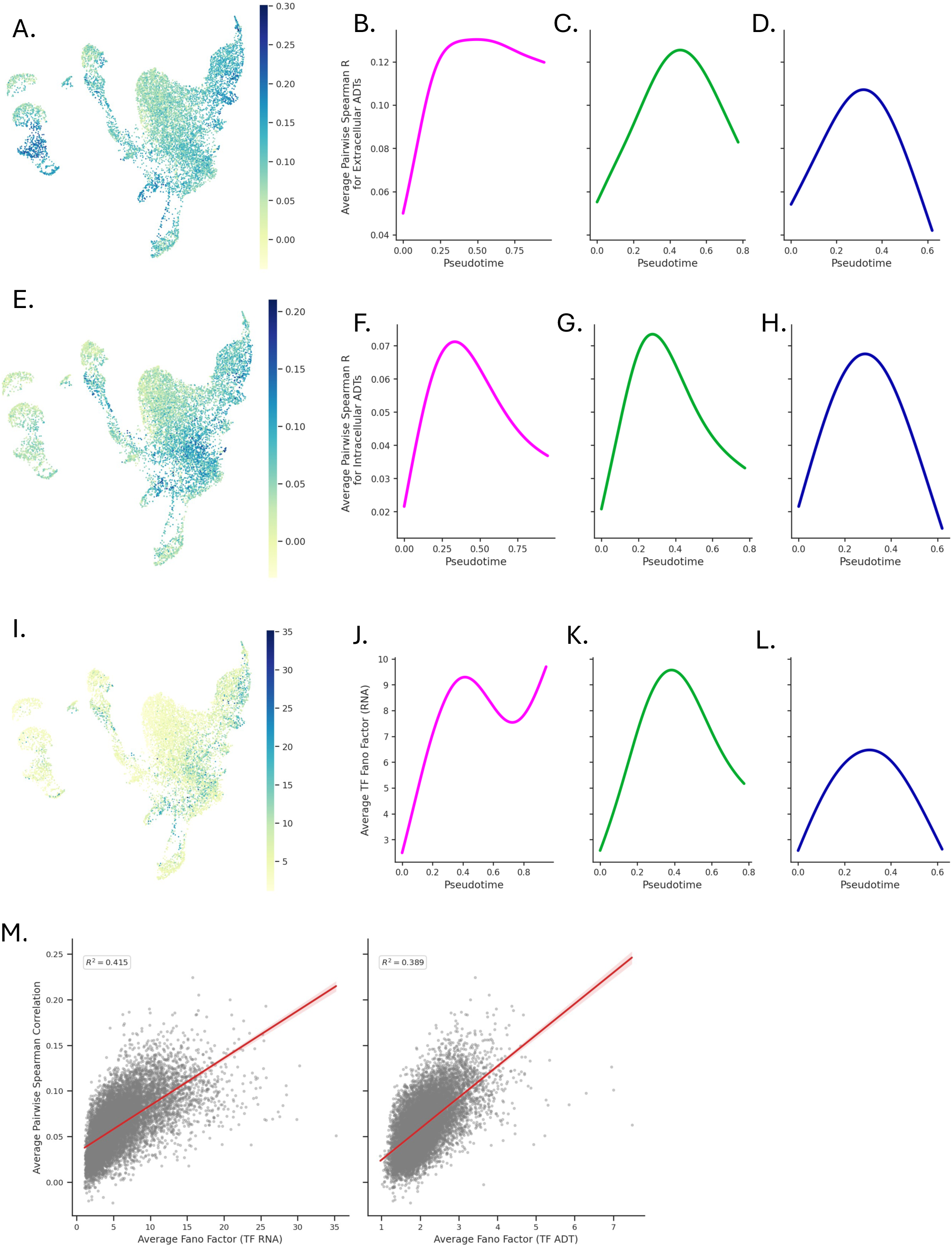
A. Average pairwise extracellular RNA-protein Spearman correlation per Milo neighborhood. **B.** Average pairwise RNA-protein Spearman correlation for extracellular ADTs along the erythroid lineage, **C.** myeloid lineage, and **D.** lymphoid lineage. **E.** Average pairwise intracellular RNA-protein Spearman correlation per Milo neighborhood. **F.** Average pairwise RNA-protein Spearman correlation for intracellular ADTs along the erythroid lineage, **G.** myeloid lineage, and **H.** lymphoid lineage**. I.** Average Fano factor of corresponding transcription factor RNAs per Milo neighborhood. **J.** Average Fano factor of transcription factor RNAs along the erythroid lineage, **K.** myeloid lineage, and **L.** lymphoid lineage. **M.** Correlation of average RNA (left) and ADT (right) transcription factor Fano factor with average protein-RNA

**Supplemental Figure 8:**
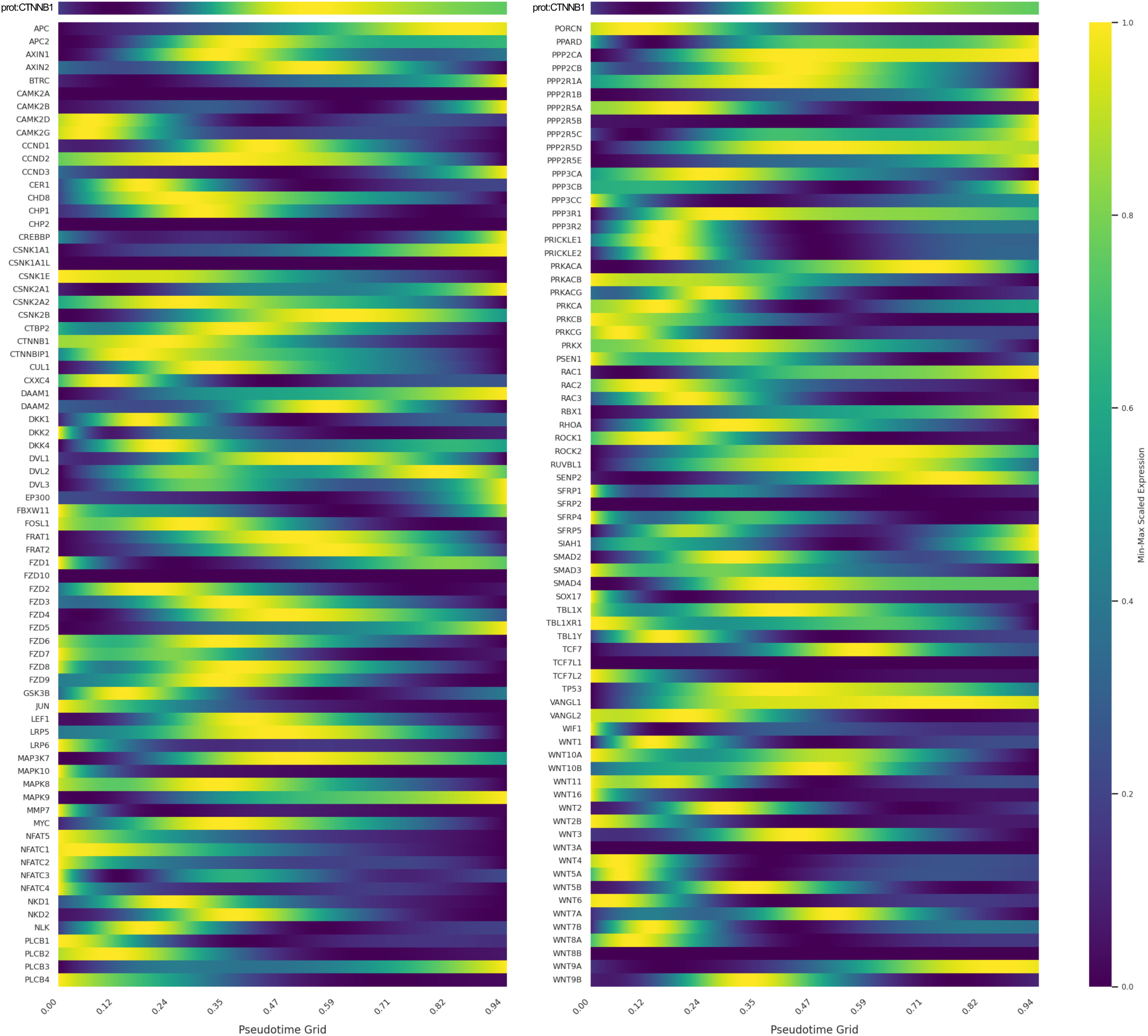
Alignment of WNT pathway genes with CTNNB1 protein expression over pseudotime. Expression of CTNNB1 protein (above, both panels) compared to expression of the 147 genes from the KEGG WNT signaling pathway found in the Flex probe set, aligned over the erythroid lineage and ordered by pseudotime.

**Supplemental Table 1:** Donor Characteristics. Table of demographics and sources per donor.

| Donor | Age | Sex | Source |
| --- | --- | --- | --- |
| Donor 1 | 23 | M | Stem Cell Technologies |
| Donor 2 | 21 | F | Cambridge Biorepository for Translational Medicine |
| Donor 3 | 33 | M | Cambridge Biorepository for Translational Medicine |
| Donor 4 | 32 | M | Stem Cell Technologies |

